# Anti-spike antibody response to natural infection with SARS-CoV-2 and its activity against emerging variants

**DOI:** 10.1101/2022.03.07.481737

**Authors:** Cheng-Pin Chen, Kuan-Ying A. Huang, Shin-Ru Shih, Yi-Chun Lin, Chien-Yu Cheng, Yhu-Chering Huang, Tzou-Yien Lin, Shu-Hsing Cheng

**Author notes:** Corresponding author: Shu-Hsing Cheng., Department of Infectious Diseases, Taoyuan General Hospital, Ministry of Health and Welfare, Taoyuan, and School of Public Health, Taipei Medical University, Taipei, Taiwan., Address: 250 Wu-Hsing St. Taipei 110, Taiwan. equal contribution (Cheng-Pin Chen, Kuan-Ying A. Huang).

## Abstract

The outbreak of severe acute respiratory syndrome coronavirus 2 (SARS-CoV-2) has substantially impacted human health globally. Spike-specific antibody response plays a major role in protection against SARS-CoV-2. Here, we demonstrated that acute SARS-CoV-2 infection elicits rapid and robust spike-binding and ACE2-blocking antibody responses, which wane approximately 11 months after infection. Serological responses were found to be correlated with the frequency of spike-specific memory B cell responses to natural infections. Further, significantly higher spike-binding, ACE2-blocking, and memory B cell responses were detected in patients with fever and pneumonia. Spike-specific antibody responses were found to be greatly affected by spike mutations in emerging variants, especially the Beta and Omicron variants. These results warrant continued surveillance of spike-specific antibody responses to natural infections and highlight the importance of maintaining functional anti-spike antibodies through immunization.

**Importance:** As spike protein-specific antibody responses play a major role in protection against SARS-CoV-2, we examined the spike-binding and ACE2-blocking antibody responses in SARS-CoV-2 infection at different time points. We found robust responses following acute infection, which waned approximately 11 months after infection. Further, the serological responses were correlated with the frequency of spike-specific memory B cell responses to natural infections. Patients with fever and pneumonia showed significantly stronger spike-binding, ACE2-blocking antibody, and memory B cell responses. Moreover, the spike-specific antibody responses were substantially affected by the emerging variants, especially the Beta and Omicron variants. These results warrant continued surveillance of spike-specific antibody responses to natural infections and highlight the importance of maintaining functional anti-spike antibodies through immunization.

## Introduction

Novel coronavirus disease (COVID-19) was first reported at the end of 2019 [1]. It spread rapidly and was declared a global health emergency [1,2]. As of February 2022, nearly 390 million confirmed cases of COVID-19 and over five million deaths have been reported to the World Health Organization [3]. The causative agent of COVID-19 is severe acute respiratory syndrome coronavirus 2 (SARS-CoV-2) [4].

SARS-CoV-2 is an enveloped beta-coronavirus with protrusions of large trimeric ‘spike’ (S) proteins. Receptor binding domains (RBDs) located at the tips of these spikes facilitate host cell entry via interaction with human angiotensin-converting enzyme 2 (ACE2) [5,6]. After entry, the SARS-CoV-2 nucleocapsid internalizes into the host cell, uses the host ribosome to produce its own mRNA, which then continuously synthesizes viral proteins in the cell cytoplasm, resulting the construction of new viral particles [7]. SARS-CoV-2 causes a broad spectrum of clinical manifestations, ranging from asymptomatic infection, mild to moderate disease including upper and lower respiratory symptoms, to critical illness requiring intubation and intensive care [1,8,9], and elicits a complex immune response [10].

Since the beginning of the pandemic, the detection and measurement of nucleoprotein (N), spike (S), and receptor-binding domain (RBD) antibodies against SARS-CoV-2 have been used to determine SARS-CoV-2 infection, outbreak investigation, seroprevalence [11,12], as well as vaccine efficacy and coverage [13,14]. Furthermore, accumulated evidence has shown the importance of antibody-mediated immunity against SARS-CoV-2 infection and the development of severe illnesses following infection in humans [15,16].

As SARS-CoV-2 continues to spread and cause outbreaks worldwide, understanding the immune response to SARS-CoV-2 is increasingly essential. Therefore, in this study, we focused on the magnitude, function, and longevity of the anti-spike antibody response to natural infection in humans. Variants of SARS-CoV-2 have been reported in many countries worldwide and harbor critical mutations in spike protein. We thus explored the impact of emerging variants on anti-spike antibodies elicited by natural infections in adults.

## Methods

### Ethics statement

The study protocol and informed consent form were approved by the ethics committee of Chang Gung Medical Foundation and Taoyuan General Hospital, Ministry of Health and Welfare. Written informed consent was obtained from each participant before inclusion in the study. The study and all associated methods were performed in accordance with the approved protocol, the principles of the Declaration of Helsinki, and Good Clinical Practice guidelines.

### Patient enrolment

Patients who were diagnosed with acute SARS-CoV-2 infection (COVID-19) by real-time reverse transcriptase-polymerase chain reaction (rRT-PCR) analysis of oropharyngeal swab samples were enrolled between January and September 2020. The patients were hospitalized in a negative-pressure isolation room according to the regulations of the Taiwan Centers for Disease Control. Blood samples were collected from the enrolled patients. The serum samples were stored at −20 °C before testing.

### Collection of respiratory samples and measurement of viral load

Briefly, 300 μL of oropharyngeal swab samples were subjected to nucleic acid extraction using the Labturbo kit (Taigen Bioscience, Taipei, Taiwan) on a LabTurbo 48 Compact System (Taigen Bioscience, Taipei, Taiwan) [17]. The isolated nucleic acids were eluted in 50 μ L elution buffer. RT-PCR was performed using the LightCycler Multiplex RNA Virus Master Kit (Roche Diagnostics, Mannheim, Germany) on a Cobas z480 analyzer (Roche Diagnostics, Mannheim, Germany), targeting envelope (E), nucleocapsid (N), and RNA-dependent RNA polymerase (RdRp) genes [18], along with an internal control. The specific primers and probes were purchased from ModularDx Kit (Tib-Molbiol, Berlin, Germany). Negative samples had cycle threshold (Ct) values higher than 37 in the reactions of both the E and RdRp genes.

### Flow cytometry

To perform a flow cytometry-based binding assay, serial dilutions of serum in 3% BSA were incubated with MDCK-RBD and MDCK-Spike cells [19] at 4 °C for 30 min. After washing, the cells were incubated with FITC-conjugated anti-human IgG secondary antibody at 4 °C for 30 min. After washing with PBS, the cells were analyzed using a BD FACSCanto II flow cytometer. At least 5,000 events were acquired for the analysis.

### Hemagglutination-inhibition assay

The RBD-binding activity of serum samples was analyzed using a hemagglutination inhibition assay in 96-well U-bottom plates. Serial dilutions of sera in PBS were mixed with VHH(IH4)-RBD [20] at room temperature. Then, a 1:40 dilution of human type O erythrocytes in PBS was added to the mixture in each well and incubated at room temperature for a further 60 min. The plate was tilted at 45 ° for at least 30 s and then examined. The endpoint was defined as the final dilution without tear drop formation.

### ACE2-blocking assay

A flow cytometry-based assay was used to analyze the RBD-ACE2 blocking activity of the serum samples. Serial dilutions of sera in PBS were mixed with biotinylated RBD [19] at room temperature. MDCK-ACE2 cells were then incubated with the mixture at 4 °C for 30 min. After washing, the ExtrAvidin-R-phycoerythrin protein was incubated with the cells at 4 °C for 30 min. After washing, the cells were analyzed using a BD FACSCanto II flow cytometer. At least 5,000 events were acquired for the analysis.

### Pseudovirus neutralization assay

HEK293T cells stably expressing human ACE2 were seeded in a 96-well plate and incubated overnight. Serial dilutions of heat-inactivated sera were prepared and mixed with pre-titrated pseudotyped lentiviruses expressing the wild-type Wuhan-1, Beta variant, Delta variant or Omicron variant spike proteins at 37 °C for 1 h. They were then inoculated to the pre-seeded cells at 37 °C, and incubated for further 16 h. The culture medium was then replaced with fresh Dulbecco’s Modified Eagle Medium (DMEM) supplemented with 1% fetal bovine serum and 100 U/mL Penicillin/Streptomycin. After a further incubation for 48 h, luciferase activity was measured using the Bright-Glo Luciferase Assay System (Promega, United States). A virus control was included for each assay. The inhibitory activity for each serum dilution was determined according to the relative light unit value as follows: [(relative light unit _Control_ - relative light unit _serum_) / relative light unit _Control_] × 100.

### Memory B cell assay

Peripheral blood mononuclear cells (PBMCs) were separated using Ficoll lymphocyte separation medium. Resuspended PBMCs were cultured in complete medium containing pokeweed mitogen (PWM), *Staphylococcus aureus* Cowan I (SAC), and CpG at 37 °C for 5 days. Cultured PBMCs were collected, washed, and resuspended for the ELISpot assay. The ELISpot assay was used to detect spike-binding IgG-, IgM-, and IgA-secreting cells. Briefly, a 96-well Millipore plate was coated with SARS-CoV-2 spike or anti-human IgG at 4 °C overnight. After washing, plates were blocked with 2% dry skim milk at 37 °C for 2 h. After washing, resuspended cultured cells were added to the wells and incubated at 37 °C for 16 h. After washing, the wells were incubated with anti-human IgG, IgM, or IgA, secondary antibodies conjugated with alkaline phosphatase at room temperature for 2 h. After washing, the wells were developed using an alkaline phosphatase substrate kit at room temperature for 2–5 minutes. The spot-forming cells were counted using an automated ELISpot plate reader [21].

### Statistical analyses

Statistical analyses were performed using GraphPad Prism or Excel. The unpaired t-test was used to determine the differences between two independent groups. One-way analysis of variance was used to determine the differences among three or more independent groups, and Tukey’s test was used for post-hoc analysis. The chi-squared test was used to analyze the relationships between categorical variables. Linear regression was used to evaluate the correlation between variables. Statistical significance was set at p < 0.05.

## Results

### The clinical manifestations and spectrum of natural SARS-CoV-2 infection

In total, 25 patients were enrolled. Their mean age was 38.68 (SD, 13.4) years. The male-to-female ratio was 12:13, and most patients were from other countries/regions. Among the subjects, five developed pneumonia, as confirmed by chest radiography or computed tomography. The mean age of patients with and without pneumonia was 47.4 and 36.5 years respectively (p = 0.105). Fever (60%) and cough (60%) were the frequently reported symptoms. Patients with pneumonia had higher level of ALT (46.8 U/L vs 26.6 U/L, p = 0.018) than those without pneumonia. Patients with and without pneumonia did not have remarkable differences in white blood cell count, C-reactive protein, creatinine, and lactate dehydrogenase, and received similar therapeutic regimens (Table 1).

**Table 1.** Demographics and clinical description of 25 COVID-19 cases

|  | Pneumonia | No<br>pneumonia | Total | P value |
| --- | --- | --- | --- | --- |
| Number (%) | 5 (20) | 20 (80) | 25(100) |  |
| Age mean (±SD) | 47.4 (10.5) | 36.5 (13.4) | 38.68 (13.4) | 0.105 |
| Male n (%) | 3 (60) | 9 (45) | 12 (48) | 0.459 |
| Female n (%) | 2 (40) | 11 (55) | 13 (52) |  |
| Imported cases n (%) | 4 (80) | 17 (75) | 21 (84) | 0.720 |
| Europe | 0 | 6 (30) | 6 (24) |  |
| America | 1 (20) | 3 (15) | 4 (16) |  |
| Africa | 1 (20) | 2 (10) | 3 (12) |  |
| Asia-pacific | 2 (40) | 6 (30) | 8 (32) |  |
| Indigenous cases n (%) | 1 (20) | 3 (15) | 4 (16) |  |
| Occupation n (%) |  |  |  | 0.720 |
| Student | 0 | 5 (25) | 5 (20) |  |
| Military | 1 (20) | 2 (10) | 3 (12) |  |
| Business | 1 (20) | 5 (25) | 6 (24) |  |
| Professional | 2 (40) | 6 (30) | 8 (32) |  |
| Others | 1 (20) | 2 (10) | 3 (12) |  |
| Subjected symptoms n (%) |  |  |  |  |
| Fever | 3 (60) | 9 (45) | 12 (48) | 0.459 |
| Cough | 3 (60) | 9 (45) | 12 (48) | 0.459 |
| Diarrhea | 1 (20) | 6 (30) | 7 (28) | 0.564 |
| Rhinorrhea | 0 | 7 (35) | 7 (28) | 0.161 |
| Sore throat | 0 | 6 (30) | 6 (24) | 0.219 |
| General soreness | 2 (40) | 4 (20) | 6 (24) | 0.343 |
| Abnormal smell | 0 | 6 (30) | 6 (24) | 0.219 |
| Abnormal taste | 0 | 5 (25) | 5 (20) | 0.292 |
| Average symptoms n (SD) | 2.4 (0.54) | 3.1 (2.28) | 2.9 (2.06) | 0.539 |
| Laboratory data |  |  |  |  |
| WBC 10<sup>9</sup>/L mean (SD) | 6252 (2653) | 5834 (1880) | 5963 (1962) | 0.865 |
| Lymphocytes 10<sup>9</sup>/L mean (SD) | 1211 (616) | 1511 (628) | 1481 (630) | 0.347 |
| CRP mg/dL mean (SD) | 1.16 (0.91) | 0.57 (0.72) | 0.67 (0.78) | 0.143 |
| Creatinine mg/dL mean (SD) | 0.96 (0.29) | 0.76 (0.13) | 0.80 (0.18) | 0.194 |
| ALT U/L mean (SD) | 46.8 (22.8) | 26.6 (13.9) | 30.6 (17.5) | 0.018 |
| LDH U/L mean (SD) | 258.8 (60.3) | 214.2 (42.4) | 224.4 (49.3) | 0.074 |
| Antiviral agent* n(%) | 4 (80) | 7 (35) | 11(44) | 0.096 |
| Immune modulator # n(%) | 4 (80) | 13 (65) | 17 (68) | 0.475 |
| Steroid n(%) | 2 (40) | 1(5) | 3 (12) | 0.091 |

\*Among pneumonia cases, 2 had lopinavir, 1 had both lopinavir and remdesivir, and 1 had oseltamivir.
Among non-pneumonia cases, 6 had oseltamivir, and 1 had lopinavir. # Hydroxychloroquine had been used in 4 pneumonia cases and 12 non-pneumonia cases. Colchicine had been used in 1 pneumonia case and 1 non-pneumonia case.

### Anti-spike antibody response

In total of 21 (84%) patients showed a detectable spike-binding antibody response in the serum at day 21 ± 8 (6–33) after the onset of illness. Patients negative for the anti-spike antibody response (n = 4) presented mild illness, and three of them had no fever (Figure 1A, Supplemental Table 1).

**Figure 1.**
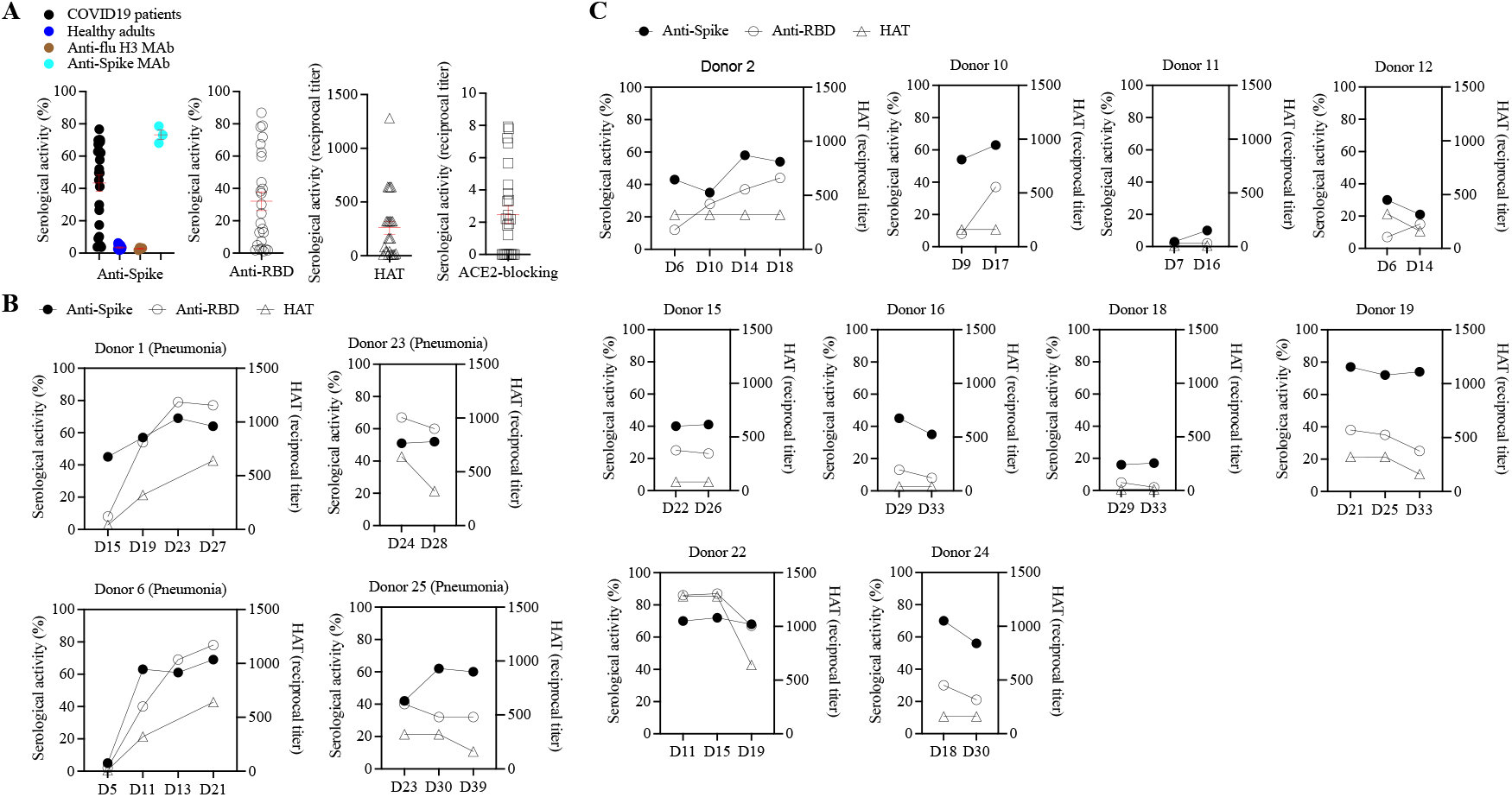
Anti-spike antibody response to natural infection with SARS-CoV-2. (A) Spike-binding, RBD-binding, haemagglutination-inhibition and ACE2-blocking activity of sera from COVID-19 adult patients (n=25). Each point represents peak serological activity for each patient and red line represents mean ± standard error of mean. Healthy adults sera collected in 2017 were included as control. Spike-binding serological activity that was above two standard deviations and the mean was positive. Anti-flu H3 and anti-SARS-CoV-2 spike human monoclonal antibodies were included as controls. (B) Magnitude and kinetics of serological anti-spike response for four COVID-19 pneumonia patients. (C) Magnitude and kinetics of serological anti-spike response for ten COVID-19 patients with mild upper respiratory tract illness. HAT, haemagglutination-inhibition titer; RBD, receptor-binding domain; D, day after onset.

The spike-binding antibody response was detected as early as the first week after onset, continued to rise between the second and third weeks, and reached a plateau at three weeks after onset during the convalescence period (Figure 1B, 1C). In most cases, the RBD-binding antibody response was detectable at the same time point, whereas the spike-binding antibodies were elicited (Figure 1).

We also examined the level of functional anti-spike antibodies in the serum using hemagglutination inhibition and ACE2-blocking assays. Nineteen (90%) patients with positive spike-binding antibody responses showed detectable hemagglutination-inhibition titers, and 15 (71%) patients showed ACE2-RBD blocking serological activities (Figure 1A).

The magnitude of the spike-binding antibody response was significantly correlated with RBD-binding, hemagglutination inhibition, and the ACE2-blocking antibody response in COVID-19 patients (Supplementary Figure 1), indicating the protective potential of anti-spike antibodies elicited upon natural infection.

## B cell response

The development of memory B cell responses to spike protein constitutes a major part of the humoral immune memory against SARS-CoV-2. Spike-specific memory B cell responses to natural infection were measured using ELISPOT (Figure 2A). Spike-specific memory B cell responses were elicited, with an average frequency of 1.3 ± 1.2% of peripheral B cells, and were detected on day 19 ± 7 (6–33) after the onset of illness. The memory B cell response was detectable within two weeks (n = 5, 0.6 ± 1.0%, day 6–13) and increased three weeks after the onset of illness (n = 15, 1.5 ± 1.2%, day 14–33), but the difference was not statistically significant (p = 0.12, unpaired two-tailed t-test). Three patients (60%) had no detectable memory B cell response in the first two weeks after onset; in contrast, one patient (7%) failed to develop a memory B cell response even at three weeks after onset. Although SARS-CoV-2 infection elicited a spike-specific B cell response, IgM memory B cells were predominantly induced, followed by IgG and IgA B cells (Figure 2B).

**Figure 2.**
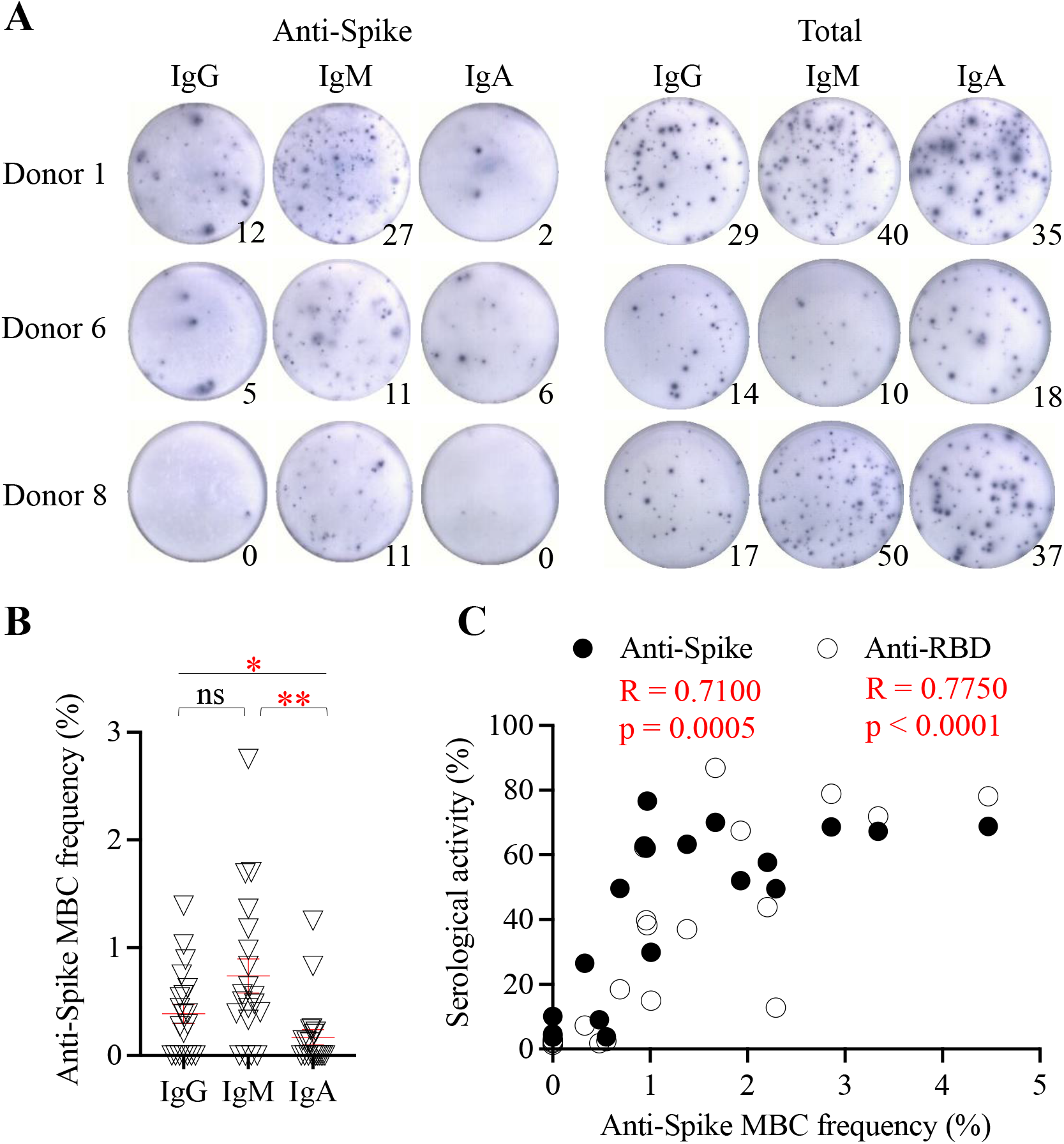
Anti-spike memory B cell response to natural infection with SARS-CoV-2. Enzyme-linked immunosorbent spot showing SARS-CoV-2 spike-specific IgG, IgM, and IgA memory B cell in the peripheral blood upon natural infection. Circulating total IgG, IgM, and IgA memory B cells were also measured. 200,000 cultured cells were added into each of anti-spike B cell wells and 5,000 cultured cells were added into each of total B cell wells. (B) Frequency of spike-specific IgG, IgM, and IgA memory B cells in the peripheral blood (n=20). Each point represents memory B cell frequency for each patient and red line represents mean ± standard error of mean. One-way ANOVA with post-hoc Tukey’s test was used to compare the difference among groups. *p = 0.05, **p = 0.01. ns, not significant. (C) Relationship of spike-specific memory B cell frequency (IgG+IgM+IgA) and spike-binding and RBD-binding serological activity among COVID-19 patients. Linear regression was used to model the relationship between two variables. MBC, memory B cell; RBD, receptor-binding domain.

Upon natural infection, the spike-specific memory B cell response was significantly correlated with the peak spike-binding and ACE2-blocking serological response, indicating a critical role of the B cell response in the development of antibody immunity upon SARS-CoV-2 infection (Figure 2C).

### Relationship of antibody response and clinical severity

Primary infection with SARS-CoV-2 can cause mild-to-severe clinical symptoms. In this study, fever duration did not correlate with the peak viral load in the respiratory sample (Supplementary Figure 2A). However, the fever duration was significantly correlated with the magnitude of spike-binding, RBD-binding antibody responses, and functional hemagglutination-inhibition titers (Supplementary Figure 2B). Patients who experienced fever had a significantly stronger RBD-binding antibody response, hemagglutination-inhibition titer, and spike-specific IgM B cell response compared with those in patients without fever (Figure 3B). Patients who developed pneumonia showed significantly stronger anti-spike antibody and B-cell responses when compared with the responses in patients without pneumonia (Figure 3A).

**Figure 3.**
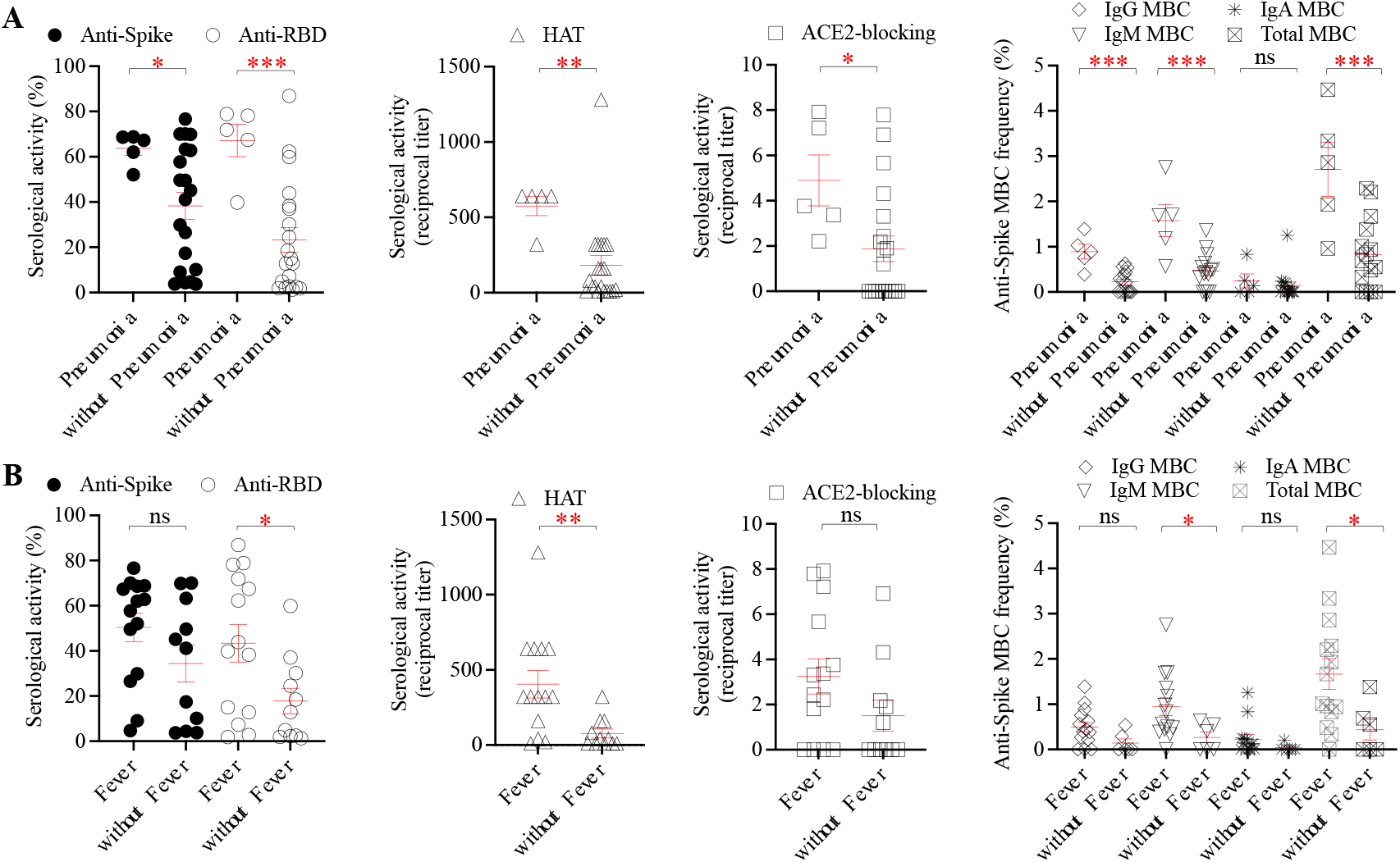
Comparison of anti-spike antibody and B cell responses between (A) patients with (n=5) and without (n=20) pneumonia and (B) patients with (n=14) and without (n=11) fever. Each point represents antibody or B cell response for each patient and red line represents mean ± standard error of mean. Unpaired two-tailed t test was used to compare the difference between two groups. *p = 0.05, **p = 0.01, ***p = 0.001. ns, not significant. RBD, receptor-binding domain; HAT, haemagglutination-inhibition titer; MBC, memory B cell.

### Longevity of the antibody response

Follow-up serum samples were collected from a subset of patients at least seven months after infection. Between enrolment and follow-up, no re-infection with SARS-CoV-2 was found among the patients. The spike-binding antibody response and functional hemagglutination-inhibition titer waned at 11 ± 3 (7–15) months after infection (Supplementary Figure 3). A substantial portion of the patients tested (75%, 6 of 8) had no detectable hemagglutination-inhibition serological titer during follow-up, and an 8- to 16-fold reduction in titer was noted in the remaining patients. An average of 2.6 ± 1.0 (1.0–3.5) fold reduction in the spike-binding antibody response was also detected, which was in accordance with the decline in the functional serological titer.

### Infection-induced anti-spike antibodies against emerging variants

The neutralization activity of convalescence and follow-up sera was then tested against the Beta, Delta, and Omicron variant pseudoviruses. The results showed that infection-induced anti-spike antibodies had greatly reduced activities against Beta, Delta, and Omicron variants in both the convalescence and follow-up sera (Figure 4). The convalescence and follow-up sera showed 83 ± 82 (15–306) and 165 ± 167 (12–456) fold reduction in the neutralization activity against the Omicron variant, respectively, suggesting a substantial impact of multiple mutations in the Omicron spike on the antibody response upon natural infection.

**Figure 4.**
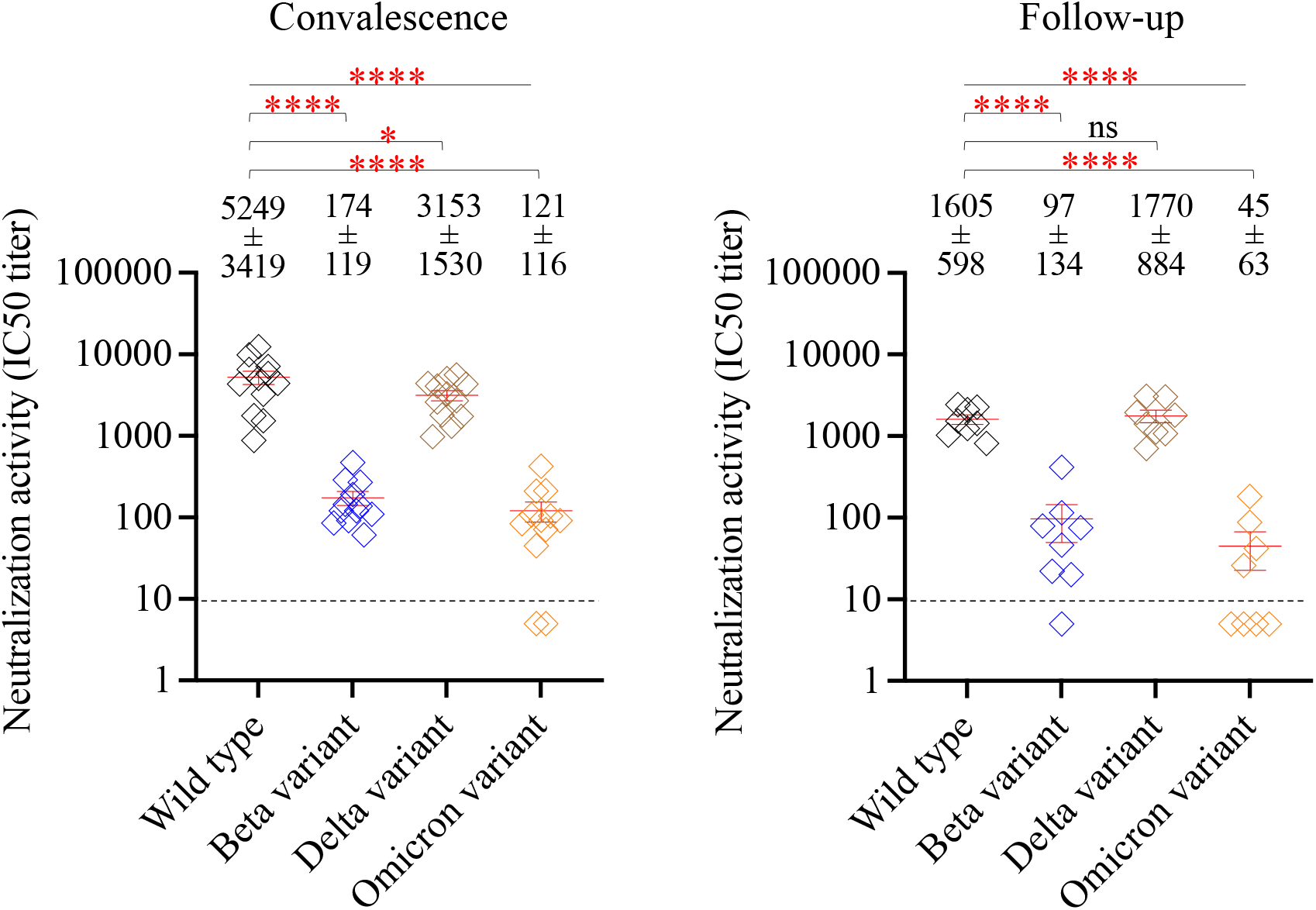
Effect of SARS-CoV-2 variants on anti-spike antibody response to natural infection. Neutralization activities of convalescene sera (n=12, left panel) and follow-up sera (n=8, right panel) agasinst wild-type, Beta variant, Delta variant and Omicron variant pseudoviruses. All sera were tested with a starting dilution of 1:10 (cut-off, dash line) and those that failed to neutralize virus at the starting dilution were recorded as IC50 titer 5. Each point represents IC50 titer for each sample and red line represents mean ± standard error of mean. The dash line represents the One-way ANOVA with post-hoc Tukey’s test was used to compare the difference among groups. *p = 0.05, ****p = 0.0001. ns, not significant. IC50, half-maximal inhibitory concentration.

## Discussion

SARS-CoV-2 belongs to the large family of coronaviruses, which includes viruses causing common cold (229E, NL63, OC43, and HKU1), Middle East respiratory syndrome coronavirus, and severe acute respiratory syndrome coronavirus. The process of antibody production is assumed to be similar to that against other seasonal coronaviruses. During SARS-CoV-2 infection, antibodies of the IgM class may be detected approximately 6 days after infection, and IgG may be detected after 8 days; concentrations of the antibodies may then decline over several months allowing subsequent infection [22,23]. In this study, we clearly demonstrated that the anti-SARS-CoV-2 spike antibody level surged as early as the first week after symptom onset, continued to rise between the second and third weeks, and reached a plateau by three weeks during the convalescence period, in line with the findings of previous studies. Antibodies that recognize the RBD have been considered the most important component of immunity against SARS-CoV-2 in humans [6,19,22,23]. In this study, the RBD-binding antibody response, with its functional activities measured by hemagglutination-inhibition and the ACE2-blocking assay, was detectable simultaneously as spike-binding antibodies were elicited. This implied that the detected anti-RBD antibodies following natural infection of wild-type SARS-CoV-2 might be neutralizing. This was further corroborated, in this study, by serological neutralizing activity against viruses bearing the wild-type and variant spike proteins.

Memory B cells circulate throughout the body in a quiescent state and are critical in an accelerated secondary immune response upon re-exposure to the pathogen [24,25]. Therefore, development of the memory B cell response to the spike protein constitutes a major part of humoral immunity against SARS-CoV-2. A previous study reported that the size of RBD-specific memory B cells may remain stable nearly six months after natural infection of SARS-CoV-2, and that these memory cells could further display clonal turnover and derived antibodies express resistance to RBD mutations [26,27]. Here, our results demonstrated that the spike-specific memory B cell response was detectable three weeks after the onset of illness, and that the memory B cell response significantly correlated with peak spike-binding and ACE2-blocking serological response, indicating a critical role of the development of B cell response. Although, the role of T cell-mediated immune response was not explored in this study, the generation of virus-specific memory B cells after SARS-CoV-2 infection could be dependent on the presence of CD4+ T helper cells [10,28].

Primary infection with SARS-CoV-2 causes mild-to-severe clinical symptoms. In previous reports, patients with severe COVID-19 may develop a stronger antibody response than those with mild illness [23,29]. Consistently, we observed that patients who experienced fever had a significantly stronger RBD-binding antibody response, hemagglutination-inhibition titer, and spike-specific IgM B cell response than those without fever. Patients who developed pneumonia showed significantly stronger anti-spike antibody and B-cell responses when compared with the responses in patients without pneumonia. Nevertheless, the magnitude of the spike-binding antibody level was more than 2-fold lower after an average of 11 months compared to that in the convalescent stage of infection. Thus, passive re-immunization through vaccination could be beneficial to boost anti-spike antibody level among individuals with prior infection.

Emerging SARS-CoV-2 variants could spread quickly, and become the dominant strains in the outbreaks [30]. For example, the Beta (B.1.351) and Delta (B.1.617.2) variants were designated as variants of concern in 2021[31]. At the end of 2021, the Omicron variant (B.1.1.529) emerged and rapidly co-circulated with the Delta variant [32,33]. Recently, the Omicron variant has become prevalent worldwide [34]. These variants have several mutations in the spike protein, with the Omicron (B.1.1.529) clade displaying over 30 changes, 15 of which are located in the RBD [35]. Further evidence indicates that spike mutations in emerging variants may enhance transmissibility and contribute to the escape of viruses from humoral immunity [36,37]. Recent research has demonstrated that convalescent sera do not provide cross-protection against new Omicron variants [38,39]. Congruously, our study demonstrated decreased neutralization activity, with more than 80- and 160-fold reductions against the Omicron variant in convalescent and follow-up sera, respectively. It has been shown that a booster immunization may elicit a prominent neutralization titer against the variant, which may reduce the risk of breakthrough infection [38,40].

Some limitations may exist in the study. First, this was a single-center observational study and the time points of blood sampling varied among patients. Second, anti-spike antibody response was undetectable in four patients. Most of them did not have risk factors, i.e., immunocompromised status, obesity, and age over 65 years, that may affect immunological responses [41,42]. However, all four patients had mild illness and one of them provided samples within the first week of illness. Third, a longitudinal follow-up of memory B cell responses was not performed. Previous research revealed that the size of antigen-specific memory B cell repertoire may show a trajectory pattern, and a long-term observation might be required to assess the dynamics of anti-spike memory B cell populations in the near future [22,23,26,27,29].

In conclusion, this study confirmed that acute SARS-CoV-2 infection elicits a rapid and robust spike-binding and ACE2-blocking antibody response, which wanes approximately 11 months after infection. Serological responses correlate with the frequency of spike-specific memory B cell responses to natural infections. Patients with fever and pneumonia develop significantly higher spike-binding, ACE2-blocking, and memory B-cell responses. However, spike-specific antibody responses are greatly affected by spike mutations in emerging variants, especially the Beta and Omicron variants. These results warrant continued surveillance of the spike-specific antibody response to natural infection and suggest the maintenance of functional anti-spike antibodies through vaccination.

## Acknowledgements

We acknowledge Alain Townsend, Tiong Kit Tan, Pramila Rijal and Lisa Schimanski for the support of VHH(IH4)-RBD. We also thank the patients and the care team of the COVID-19 designated wards at Taoyuan General Hospital, Ministry of Health and Welfare.

## Author contributions

Conceptualization: Cheng-Pin Chen, Kuan-Ying A. Huang, Shu-Hsing Cheng Cheng-Pin Chen, Kuan-Ying A. Huang, Shin-Ru Shih, Yi-Chun Lin, Chien-Yu Cheng, Yhu-Chering Huang, Tzou-Yien Lin and Shu-Hsing Cheng

Data curation: Cheng-Pin Chen, Kuan-Ying A. Huang, Yi-Chun Lin, Chien-Yu Cheng and Shu-Hsing Cheng

Formal analysis: Kuan-Ying A. Huang, Shin-Ru Shih, Shu-Hsing Cheng

Funding Acquisition: Kuan-Ying A. Huang

Investigation: Cheng-Pin Chen, Kuan-Ying A. Huang, Shu-Hsing Cheng

Methodology: Kuan-Ying A. Huang, Shin-Ru Shih

Project administration: Kuan-Ying A. Huang, Shu-Hsing Cheng

Resources: Kuan-Ying A. Huang, Shu-Hsing Cheng

Software: Kuan-Ying A. Huang

Supervision: Yhu-Chering Huang, Tzou-Yien Lin

Validation: Cheng-Pin Chen, Kuan-Ying A. Huang, Shu-Hsing Cheng

Visualization: Cheng-Pin Chen, Kuan-Ying A. Huang

Writing-original draft: Cheng-Pin Chen, Kuan-Ying A. Huang, and Shu-Hsing Cheng

Writing-review and editing: Cheng-Pin Chen, Kuan-Ying A. Huang, Shin-Ru Shih, Yi-Chun Lin, Chien-Yu Cheng, Yhu-Chering Huang, Tzou-Yien Lin, and Shu-Hsing Cheng

