## Supplemental Table1 for "Anti-spike antibody response to natural infection with SARS-CoV-2 and its activity against emerging variants"

### Supplemental Table 1. Demographic and clinical manifestations of 25 COVID-19 patients.

| Donor | Gender | Age | Symptoms and signs | Fever duration (Day) | Clinical diagnosis | Samples taken (Day after onset) |
|---|---|---|---|---|---|---|
| 1 | F | 55 | Fever, cough | 12 | Pneumonia | D15, 19, 23, 27 |
| 2 | M | 52 | Fever | 5 | Febrile illness | D6, 10, 14, 18 |
| 3 | M | 30 | Cough | 0 | URI | D13 |
| 4 | F | 26 | Fever, cough | 3 | URI | D10 |
| 5 | F | 23 | Fever, cough | 0 | URI | D9 |
| 6 | M | 43 | Fever, cough | 7 | Pneumonia | D5, 11, 13, 21 |
| 7 | F | 28 | Fever | 4 | Febrile illness | D11 |
| 8 | F | 22 | Rhinorrhea | 0 | URI | D18 |
| 9 | F | 21 | Fever, rhinorrhea | 2 | URI | D6 |
| 10 | F | 63 | Cough | 0 | URI | D9, 17 |
| 11 | M | 41 | Diarrhea | 0 | Viral syndrome | D7, 16 |
| 12 | M | 32 | Fever, cough | 1 | URI | D6, 14 |
| 13 | F | 43 | Cough | 0 | URI | D33 |
| 14 | M | 31 | Fever, cough | 8 | Pneumonia | D25 |
| 15 | M | 43 | Cough | 0 | URI | D22, 26 |
| 16 | F | 22 | Sorethroat | 0 | URI | D29, 33 |
| 17 | F | 36 | Fever, cough | 4 | URI | D21 |
| 18 | M | 41 | Sorethroat | 0 | URI | D29, 33 |
| 19 | M | 20 | Fever, sorethroat | 4 | URI | D21, 25, 33 |
| 20 | M | 58 | Fever, cough | 0 | URI | D26 |
| 21 | M | 46 | Fever, cough | 3 | URI | D24 |
| 22 | M | 55 | Fever, cough | 1 | URI | D11, 15, 19 |
| 23 | F | 56 | Fever, cough | 10 | Pneumonia | D24, 28 |
| 24 | F | 28 | Nausea | 0 | Viral syndrome | D18, 30 |
| 25 | M | 52 | Fever, cough | 12 | Pneumonia | D23, 30, 39 |

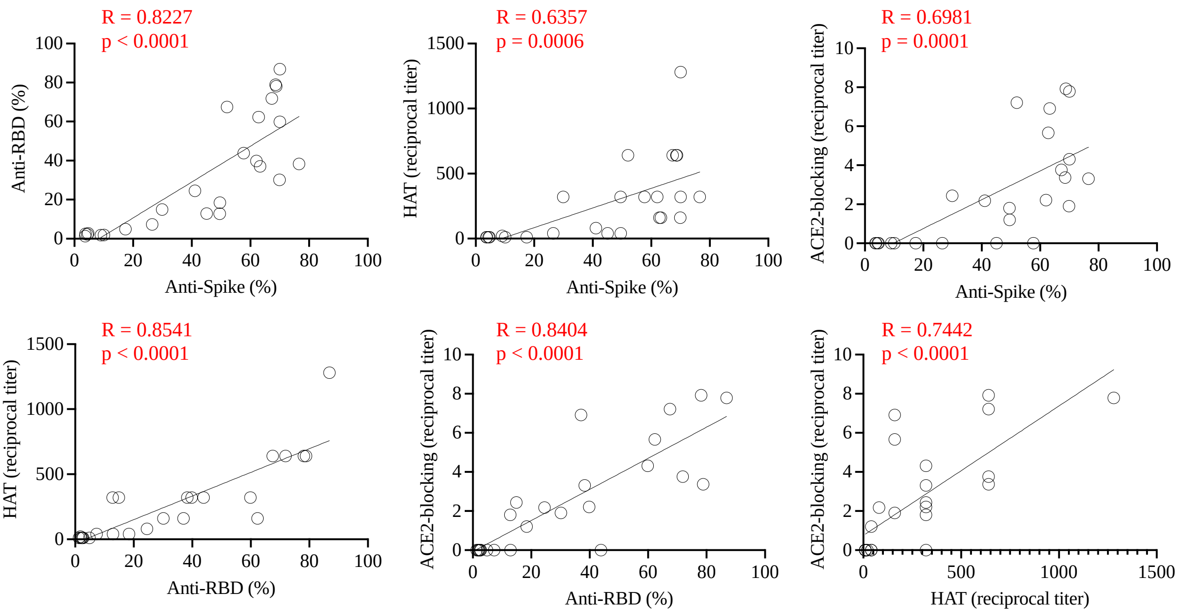


**Supplemental Figure 1. Relationship of serological spike-binding, RBD-binding, haemagglutination-inhibition and ACE2-blocking activity among COVID-19 patients.** Lnear regression is used to model the relationship between two variables.

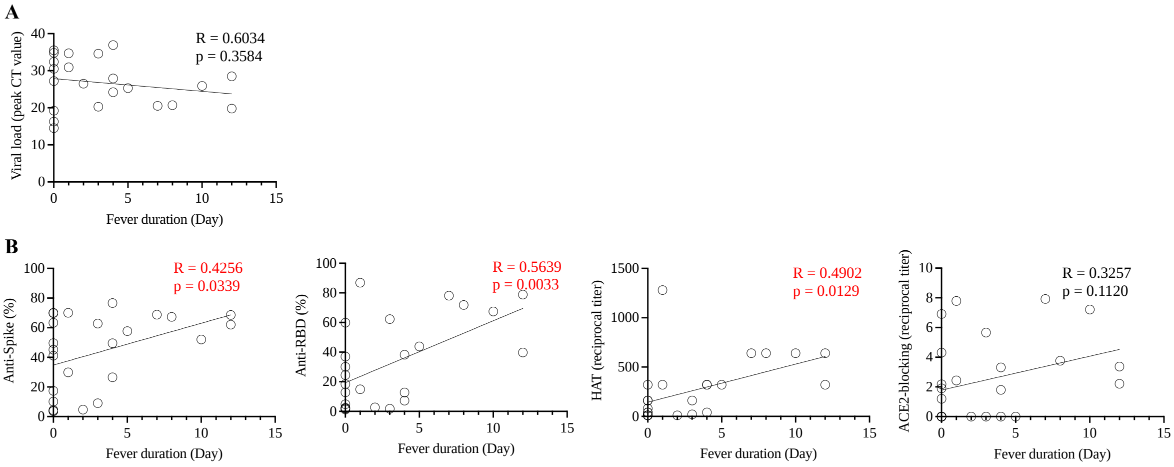


**Supplemental Figure 2.** (**A**) Relationship of fever duration and peak viral load among COVID-19 patients. (**B**) Relationship of fever duration and RBD-binding, haemagglutination-inhibition and ACE2-blocking activity among COVID-19 patients. Linear regression was used to model the relationship between two variables.

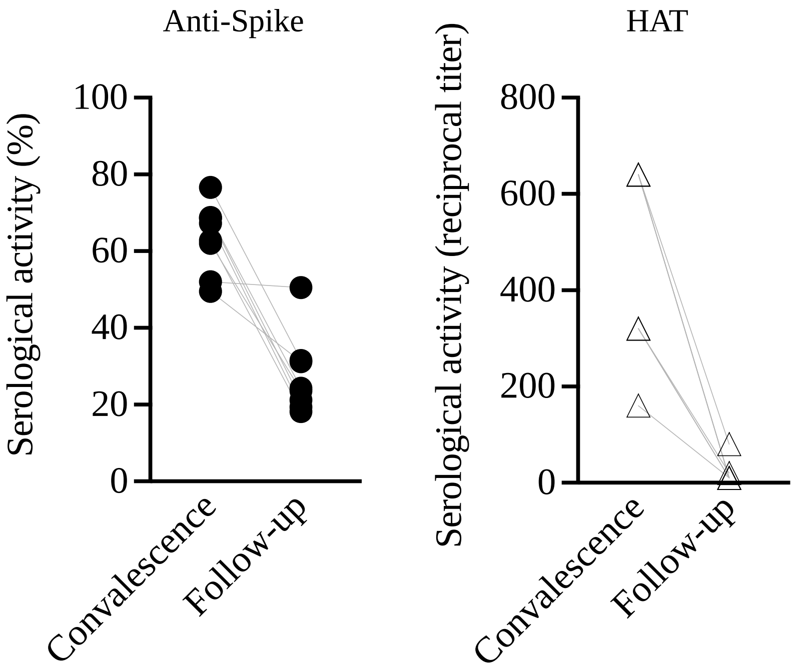


**Supplemental Figure 3.** Analysis of spike-binding antibody response and haemagglutination-inhibition serological titer at enrolment and follow-up (n=8).
